# HexSDF are Required for Synthesis of a Novel Glycolipid that Mediates Daptomycin and Bacitracin Resistance in *C. difficile*

**DOI:** 10.1101/2022.12.06.519413

**Authors:** Anthony G. Pannullo, Ziqiang Guan, Howard Goldfine, Craig D. Ellermeier

**Affiliations:** Department of Microbiology and Immunology, Carver College of Medicine, University of Iowa, Iowa City, IA 52242; Department of Biochemistry, Duke University Medical Center, Durham, NC 27710; Department of Microbiology, Perelman School of Medicine, University of Pennsylvania, Philadelphia, PA 19104; Graduate Program in Genetics, University of Iowa, Iowa City, IA 52242, USA

**Keywords:** Two-component regulatory system, cell envelope, signal transduction, gene expression, Glycolipid synthesis, membrane biogenesis

## Abstract

*Clostridioides difficile* is a Gram-positive opportunistic pathogen that results in 250,000 infections, 12,000 deaths, and $1 billion in medical costs in the US each year. There has been recent interest in using a daptomycin analog, Surotomycin, to treat *C. difficile* infections. Daptomycin interacts with both phosphatidylglycerol and Lipid II to disrupt the membrane and halt peptidoglycan synthesis. *C. difficile* has an unusual lipid membrane composition as it has no phosphatidylserine or phosphatidylethanolamine, and ∼50% of its membrane is composed of glycolipids, including the unique *C. difficile* lipid aminohexosyl-hexosyldiradylglycerol (HNHDRG). We identified a two-component system (TCS) HexRK that is required for *C. difficile* resistance to daptomycin. Using RNAseq we found that HexRK regulates a three gene operon of unknown function *hexSDF*. Based on bioinformatic predictions, *hexS* encodes a monogalactosyldiacylglycerol synthase, *hexD* encodes a polysaccharide deacetylase, and *hexF* encodes an MprF-like flippase. We find that deletion of *hexRK* leads to a 4-fold decrease in daptomycin MIC, and that deletion of *hexSDF* leads to an 8-16-fold decrease in daptomycin MIC. The Δ*hexSDF* mutant is also 4-fold less resistant to bacitracin but no other cell wall active antibiotics. Our data indicate that in the absence of HexSDF the phospholipid membrane composition is altered. In WT *C. difficile* the unique glycolipid, HNHDRG makes up ∼17% of the lipids in the membrane. However, in a Δ*hexSDF* mutant, HNHDRG is completely absent. While it is unclear how HNHDRG contributes daptomycin resistance, the requirement for bacitracin resistance suggests it has a general role in cell membrane biogenesis.

**Importance:** *Clostridioides difficile* is a major cause of hospital acquired diarrhea and represents an urgent concern due to the prevalence of antibiotic resistance and the rate of recurrent infections. Little is understood about *C. difficile* membrane lipids, but a unique glycolipid, HNHDRG, has been previously identified in *C. difficile* and, currently, has not been identified in other organisms. Here we show that HexSDF and HexRK are required for synthesis of HNHDRG, and that production of HNHDRG impacts resistance to daptomycin and bacitracin.

## Introduction

*Clostridioides difficile* a Gram-positive, obligate anaerobe, opportunistic pathogen. *C. difficile* is a common nosocomial disease that infects approximately 220,000 and killing about 12,000 people each year in the United States (Centers for Disease Control and Prevention (U.S.), 2019). *C. difficile* infections are estimated to lead to almost $1 billion in excess medical costs annually in the United States (Centers for Disease Control and Prevention (U.S.), 2019). *C. difficile* infections often occur due to disruption of the microflora caused by antibiotic treatment (Kelly et al., 2010; Slimings & Riley, 2014). A healthy microbiota helps prevent *C. difficile* colonization and subsequent infection. Evidence suggests that antibiotic treatment disrupts the microbiome which leads to changes in the production of bile salts to which *C. difficile* is sensitive (Giel et al., 2010; Koenigsknecht et al., 2015; Sorg & Sonenshein, 2008; Theriot et al., 2014, 2016; Wilson, 1983). These changes are correlated with resistance to *C. difficile* infection (Buffie et al., 2015; Theriot et al., 2016). However, competition for nutrients may also play an important role. *C. difficile* can utilize amino acids as a sole carbon and energy source using Stickland metabolism where one amino acid is an electron donor while another amino acid is an electron acceptor (Bouillaut et al., 2013, 2015). Recent evidence has found a correlation with nutrient limitation, caused by species in the microbiota that perform Stickland metabolism, may be a factor involved in microbiota-mediated inhibition of *C. difficile* colonization (Aguirre et al., 2021). Treatment of *C. difficile* infections often requires antibiotic treatments which can lead to further disruption of the microbiome and can predispose individuals to recurrent *C. difficile* infections (Bauer et al., 2011; Cornely et al., 2012; Hopkins & Wilson, 2018).

Surotomycin, a daptomycin derivative, was developed as a treatment option for *C. difficile* and its efficacy to treat *C. difficile* infections was the subject of Phase III clinical trials. The results of trials showed conflicting interpretations, one stating that Surotomycin displayed equivalent or better efficacy in treating a CDI compared to vancomycin, and the other stated that Surotomycin did not display any improvement over vancomycin (Boix et al., 2017; Daley et al., 2017; Knight-Connoni et al., 2016). Daptomycin is a cyclic lipopeptide that is used to treat antibiotic resistance strains of *Enterococcus faecalis* and *Staphylococcus aureus* (Hanberger et al., 1991; Louie et al., 1993; Snydman et al., 2000; Tally et al., 1999). Daptomycin was long thought to exert antimicrobial activity by depolarization of the membrane (Allen et al., 1987; Silverman et al., 2003), however, recent evidence suggests the mechanism may be more complicated. Daptomycin can form a complex with the peptidoglycan biosynthesis intermediates Lipid II, UPP, and UP in the presence of phosphatidylglycerol, and this complex is thought to inhibit PG biosynthesis (Grein et al., 2020; Müller et al., 2016). Daptomycin also activates the LiaRS two-component regulatory system in *B. subtilis* which is activated by other Lipid II binding antibiotics (Hachmann et al., 2009; Jordan et al., 2006; Mascher et al., 2004). Since daptomycin affects components of both the cell wall and lipid membrane, it can serve as useful tool to dissect cell envelope biogenesis.

The lipid membranes of bacteria are an essential component of the cell envelope. The lipid membrane acts as an addition barrier to the environment, generates the proton motive force, is utilized for synthesis of ATP, and sequesters nutrients. Since the membrane is integral to a myriad of cellular processes, proper synthesis and maintenance of the membrane is critical for survival. While membrane lipid synthesis has been studied in model organisms such as *E. coli* and *B. subtilis*, it is unclear if these findings translate to less studied organisms such as *C. difficile. E. coli* generally has a membrane composition of ∼70% phosphatidylethanolamine (PE), 25% phosphatidylglycerol, and 5% cardiolipin (López-Lara & Geiger, 2017; Raetz & Dowhan, 1990). *B. subtilis* has a membrane composition of ∼70% phosphatidylglycerol, ∼12% PE, ∼5% cardiolipin, ∼2% lysophosphatidylglycerol, and ∼8% glycolipids (Clejan et al., 1986).

The polar membrane lipids of *C. difficile* have not been well studied, and the polar lipid composition was only recently determined (Guan et al., 2014). The *C. difficile* membrane contains ∼30% phosphatidylglycerol, 16% cardiolipin and 50% glycolipids (Guan et al., 2014). Notably, amongst the phospholipids in *C. difficile* no PE, phosphatidylserine (PS) or lysophosphatidylglycerol has been identified (Guan et al., 2014). There are several glycolipids that make up the lipidome of the *C. difficile* membrane including mono-hexose-diradylglycerol (MHDRG) (14%), Di-hexose-diradylglycerol (DHDRG) (15%), Tri-hexose-diradylglycerol (THDRG) (5%), and a novel glycolipid, aminohexosyl-hexosyldiradylglycerol (HNHDRG) (16%) (Guan et al., 2014). In *C. difficile* the phospholipids cardiolipin and phosphatidylglycerol contained significant amounts of plasmalogens (Guan et al., 2014). Plasmalogen species are lipids where one of the ester bonds has been substituted with an ether (Goldfine, 2010; Řezanka et al., 2012), thus we refer to the lipids as diradylglycerol rather than diacylglycerol. The specific sugar(s) on these glycolipids are not known thus we refer to them has hexosyl-diradylglycerol. While MHDRG and DHDRG have been identified in other bacteria (Brundish et al., 1966; Clejan et al., 1986; Shaw, 1975), it is important to note that HNHDRG has to date only been identified in *C. difficile* (Guan et al., 2014). Finally,

Two-component systems (TCS) are a large family of regulatory systems that can be found in many forms of life including many prokaryotes and some eukaryotes, particularly yeasts and plants (Stock et al., 2000). The function of TCS can be quite varied, and in the case of many pathogens, TCS often regulate virulence factors and drug resistance mechanisms (Arthur & Quintiliani, 2001; Fournier et al., 2000; Giraudo et al., 1999; Groisman et al., 1989; Miller et al., 1989; Stephenson & Hoch, 2002). *C. difficile* is predicted to have approximately 50 TCS, with slight variations being observed between strains (Karp et al., 2019). However, the vast majority of these TCS remain unstudied, with only a handful of TCS having any identified function or signal. The *cprABCK* locus has been found to respond to a subset of lantibiotic compounds, and regulate the expression of lantibiotic resistance mechanisms, mainly in the form of an ABC transporter (McBride & Sonenshein, 2011; Suarez et al., 2013). This system shares a similar mechanism to the bacitracin resistance TCS in *Bacillus subtills, bceRSAB* (Bernard et al., 2007; Ohki et al., 2003). The response regulator Spo0A is a major regulator of sporulation, however its cognate histidine kinase has not been identified, and the manner in which Spo0A becomes activated is still unclear (Deakin et al., 2012; Mackin et al., 2013; Rosenbusch et al., 2012). The WalRK TCS is essential and required for proper cell wall morphology in *C. difficile* (Müh et al., 2022). The genes encoding the CmrST TCS, are phase variable and consists of two RRs and a single HK. CmrRST regulates a variety of cell functions, including colony morphology and motility (Garrett et al., 2019, 2022). The *agrACDB* locus, homologous to the *agr* locus in *S. aureus*, modulates virulence by increasing production of flagella and TcdA (Ahmed et al., 2020; Choudhary et al., 2018; Martin et al., 2013). The RR CD1688 has been shown to be a repressor of sporulation that functions by inhibiting expression of *spoIIR*, an important early-stage regulator of sporulation (Kempher et al., 2022). Loss of CD1688 leads to increased sporulation (Kempher et al., 2022). Understanding the roles of these unstudied TCS can give a better understanding of how *C. difficile* responds to its environment, particularly the complex host environment.

Here we describe the identification of the TCS, *hexRK*, and its regulon, *hexSDF*. We show that HexRK and HexSDF mediate daptomycin and bacitracin resistance in *C. difficile*. We also show that HexSDF is required for synthesis of the novel glycolipid HNHDRG, and that loss of HexSDF leads to large changes in lipid content of the cell membrane.

## Results

### Identification of *hexRK* and *hexSDF*

We performed a Tn-seq experiment in which sought to identify genes that are required for growth in the presence of daptomycin. We generated a transposon insertion library containing ∼80,000 colonies using pRPF215 as previously described (Dembek et al., 2015). The transposon library was then exposed to 0 and 2 μg/mL daptomycin. Upon sequencing we found there were ∼45,000 unique transposon insertions which was lower than expected. However, we identified one gene of interest *cdr202091_2610* (hereafter referred to as *hexK*) which had less insertions recovered when cells were grown with daptomycin (Table S1). *hexK* is a predicted to encode a TCS sensor histidine kinase (HK) with 2 transmembrane domains and an approximately 120 amino acid extracytoplasmic domain as predicted by DeepTMHMM (Hallgren et al., 2022). *hexK* is immediately downstream of *cdr20291_2611* (hereafter referred to as *hexR*), a predicted TCS RR. Through bioinformatic analysis we found that HexK is homologous to the relatively uncharacterized HK from *B. subtilis*, YrkQ (Supplemental Figure 1). Due to our interest in signal transduction, we focused on HexRK for survival in the presence of daptomycin.

To confirm that loss of *hexRK* led to a decrease in daptomycin resistance, as the Tn-seq data suggested we utilized CRISPRi to knockdown expression of the *hexRK* operon. We found knockdown of *hexR* lead to ∼4-fold decrease in daptomycin resistance (Figure 1B). We then constructed a Δ*hexRK* deletion using CRISPR mutagenesis and we found that loss of HexRK decreased daptomycin resistance ∼4-fold (Figure 1C). We then asked if we could restore daptomycin resistance by expression of HexR^D56E^ (a mutant of the putative phosphorylation site), which mimics phosphorylation of HexR, and likely increases HexR activity. When *hexR*^*D56E*^ is expressed from a xylose-inducible promoter in a *ΔhexRK* mutant we observed an increase in daptomycin resistance to near wild type levels (Figure 1D). Taken together our data suggests that HexRK are required for daptomycin resistance.

**Figure 1.**
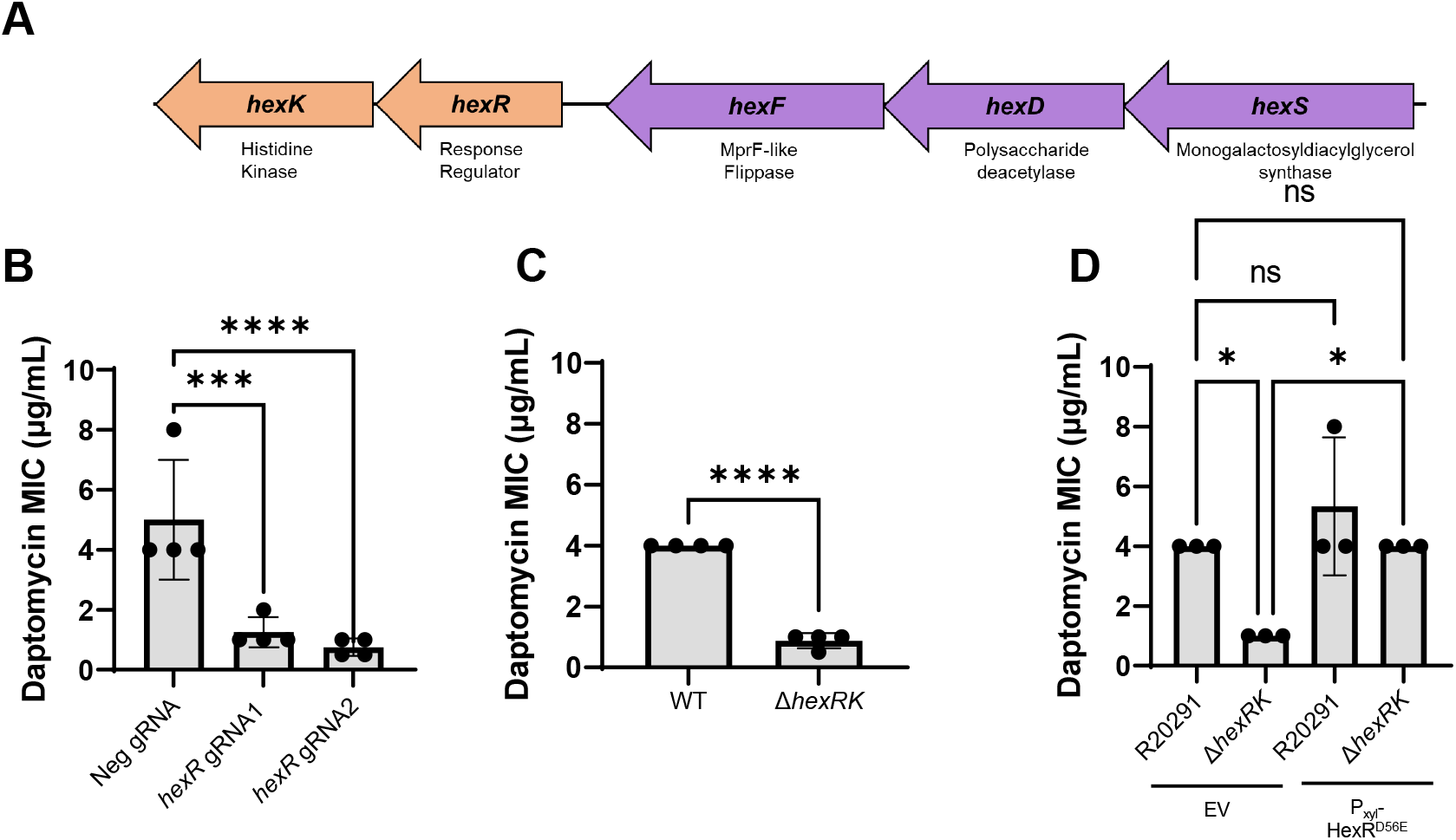
*hexRK* and *hexSDF*. (A) Diagram of *hexRK* and *hexSDF* and their respective annotations. (B) CRISPRi knockdown with guides targeting *hexR* result in ∼4-fold decreases in daptomycin resistance. (C) Δ*hexRK* has ∼4-fold lower daptomycin MIC compared to WT. Data were analyzed with an unpaired t-test, ****, *P*<0.0001. (D) Overproduction of HexR^D56E^ in Δ*hexRK* restores daptomycin resistance to WT levels. Unless otherwise specified, data were analyzed using a one-way analysis of variance with Dunnett’s multiple comparison test. ****, *P*<0.0001; *, P*<0*.*05*; NS, *P*>0.05.

### Identification of the HexR Regulon

We hypothesized that HexRK likely regulates expression of genes involved in daptomycin resistance. Thus, we performed RNA-seq, comparing WT with Δ*hexRK* to identify any genes that were differentially regulated in the absence of *hexRK*. Previous RNA-seq experiments showed *hexRK* was expressed at relatively high levels in log phase growth (Müh et al., 2022) thus we performed RNA-seq experiments on cultures grown to mid-log (OD_600_ 0.7-0.8)(Table S2).

We found 32 genes, not including *hexRK*, were differentially regulated between WT and Δ*hexRK* with a *P* value < 0.05 and a Log2 fold change 2> or <-2. Of these 32 genes in WT vs Δ*hexRK* 24 showed an increase in expression and 8 showed a decrease in expression (Table 1, Figure 2A). Since our data suggests that HexRK is required for daptomycin resistance we were most interested in those genes whose expression was dependent upon HexRK. We were particularly interested in a three gene operon *cdr20291_2614-2613-2612* (*hexSDF*) located immediately upstream of *hexRK* (Figure 1A). The genes in the *hexSDF* operon were expressed ∼12-fold lower in the Δ*hexRK* mutant (Table 1, Figure 2A). The RNA-seq revealed other differentially regulated genes, however several of these genes are reported to be phase-variable, including *cdr0440, cdr0962*, and *cdr0963, cdr3418*, and *cdr1514* (Sekulovic et al., 2018). As such, we chose to focus our efforts on better understanding the function of the *hexSDF* operon. To confirm that HexRK are required for *hexSDF* expression we constructed a P_*hexS*_*-sLuc*^*opt*^ reporter. We found that expression of the P_*hexS*_*-sLuc*^*opt*^ reporter was reduced by 26-fold in Δ*hexRK* when compared to WT, indicating that expression of *hexSDF* is dependent upon HexRK (Figure 2B). Since we found that daptomycin resistance decreases in the absence of HexRK we were interested in if daptomycin was an activator of the HexRK system. We tested if the P_*hexS*_*-sLuc*^*opt*^ was induced after a 2-hour treatment with varying concentrations of daptomycin. We found under these conditions that daptomycin (Figure 2C) did not lead to increased expression of the *hexSDF* locus.

**Table 1.**
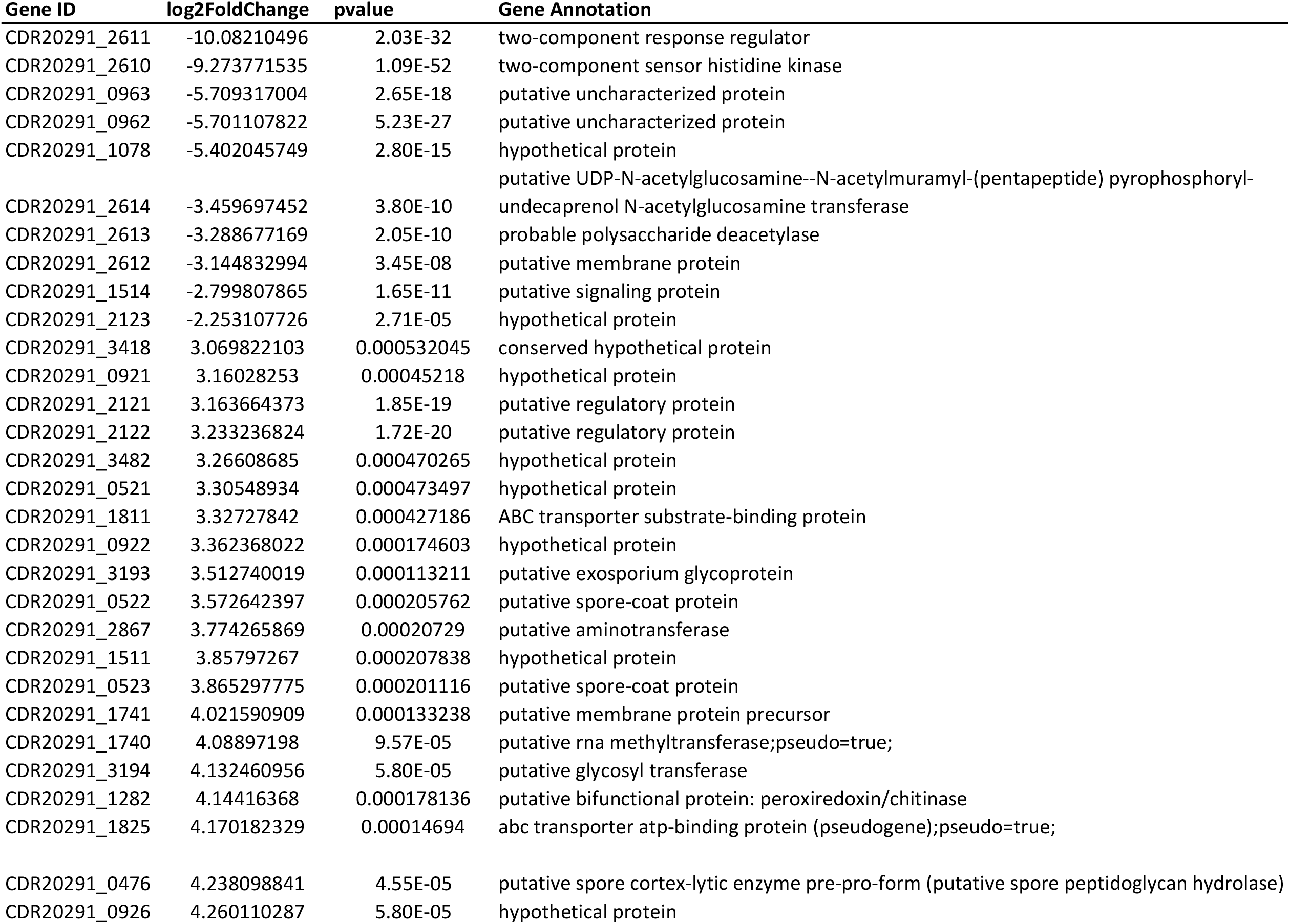
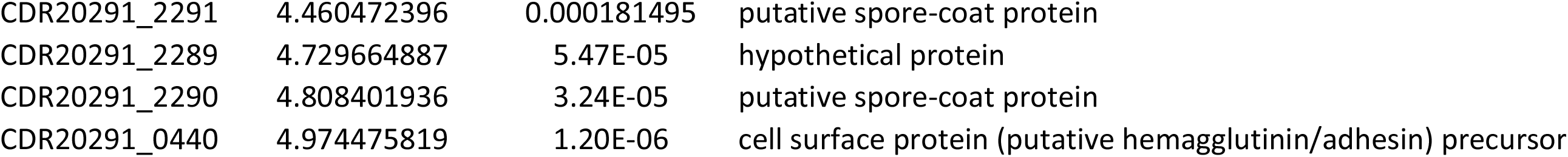
- WT v Δ*hexRK* RNA-seq Genes of Interest.

**Figure 2.**
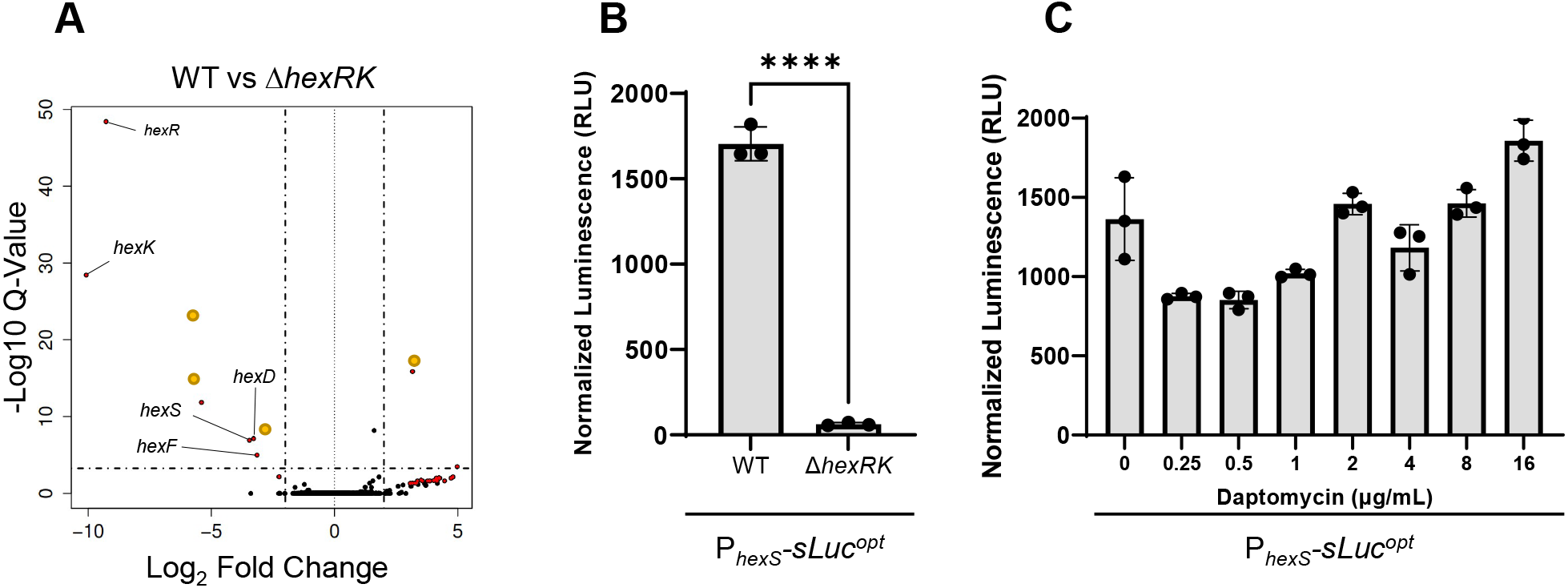
Expression of *hexSDF* is regulated by HexRK. (A) Volcano plot of RNA-seq data comparing WT and Δ*hexRK* shows a decrease in the expression *hexSDF*, and relatively few other changes in the transcriptome. Genes that have been previously reported to be phase variable have been highlighted with a yellow dot. (B) The luciferase reporter, P_*hexS*_-*sLuc*^*opt*^, shows significantly decreased activity in Δ*hexRK* comparted to WT, confirming the differential expression identified in RNA-seq. Data were analyzed with an unpaired t-test, ****, *P>*0.0001. (C) The P_*hexS*_-*sLuc*^*opt*^ reporter does not show altered luminescence in the presence of multiple concentrations of daptomycin after incubation for 2 hours.

### Loss of *hexRK* and *hexSDF* Leads to Daptomycin and Bacitracin Susceptibility

HexS is annotated as a putative monogalactosyldiacylglycerol synthase, HexD is a predicted polysaccharide deacetylase, and HexF is annotated as MprF-like. In other organisms MprF is a known to contribute to daptomycin resistance (Figure 1A) (Ernst et al., 2018; Peschel et al., 2001; Pillai et al., 2007). MprF from *S. aureus* and *B. subtilis* is a bi-functional protein that adds lysine onto phosphatidylglycerol generating Lys-PG and then flips Lys-PG from the inner leaflet of the membrane to the outer leaflet (Ernst et al., 2009; Salzberg & Helmann, 2008; Staubitz et al., 2004). Bioinformatic analysis reveals that HexF contains homology to the flippase domain of MprF, but HexF does not appear to contain the domain required for lipid lysinylation (Supplemental Figure 2). Based on the bioinformatic predictions of the *hexSDF* operon, we hypothesized that the operon likely encodes proteins that alter biosynthesis of membrane lipids.

To determine if HexSDF are required for daptomycin resistance we utilized CRISPRi to knockdown expression of the *hexSDF* operon. We found that knockdown of *hexSDF* led to ∼4-fold decrease in daptomycin MIC (Figure 3A). Having demonstrated CRISPRi knockdowns of *hexSDF* decreased daptomycin resistance we sought to construct a deletion of these genes to determine if they are necessary for daptomycin resistance. We constructed a deletion of the *hexSDF* operon (Δ*hexSDF*) and found that this resulted in an ∼8-fold decrease in daptomycin resistance compared to WT (Figure 3B). We then complemented these mutants by expressing *hexSDF* from a xylose inducible promoter. We found that overproduction of HexSDF in WT R20291 did not lead to significant increases in daptomycin resistance, however, when HexSDF is overproduced in either *ΔhexRK* or Δ*hexSDF*, daptomycin resistance is restored to WT levels (Figure 3C). Taken together this data suggests that HexSDF is required for HexRK-mediated daptomycin resistance.

**Figure 3.**
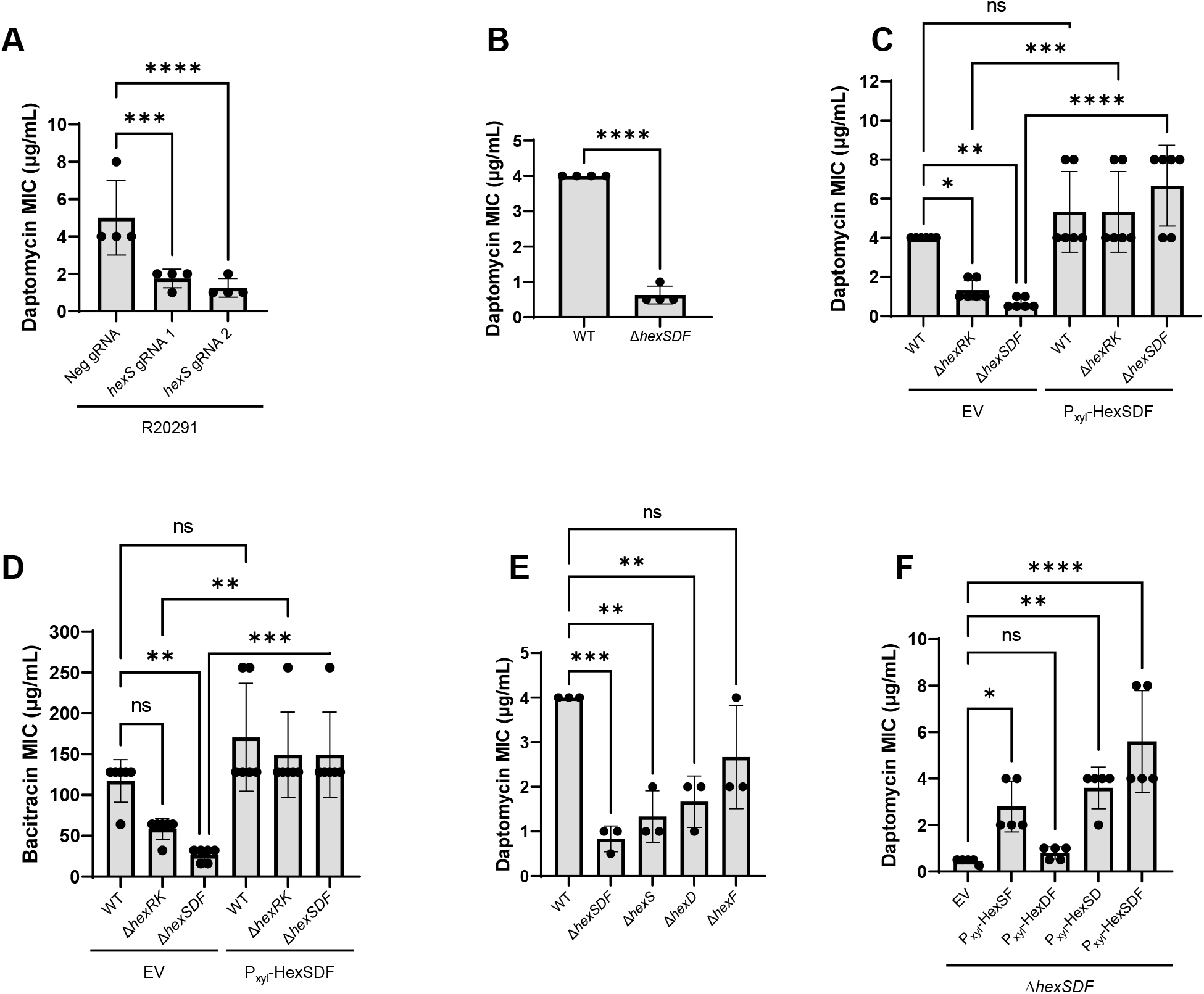
HexRK and HexSDF are involved in daptomycin and bacitracin resistance. (A) CRISPRi knockdown of *hexS* leads to ∼4-fold decrease in daptomycin resistance (B) Deletion of *hexSDF* results in ∼8-fold decrease in daptomycin MIC. Data were analyzed using unpaired t-test, ****, *P*<0.0001. (C) Overproduction of HexSDF restores daptomycin resistance in Δ*hexSDF*. (D) Deletion of *hexSDF* results in ∼4-fold decrease in bacitracin MIC, and overproduction of HexSDF restores bacitracin resistance in Δ*hexSDF*. (E) Deletion of *hexS* or *hexD* decreases daptomycin resistance. (F) Overproduction of HexSD, HexSF, and HexSDF lead to increases in daptomycin MIC in Δ*hexSDF*. Unless otherwise specified, data were analyzed using one-way analysis of variance using Dunnett’s multiple comparisons test. ****, P<0.0001; ***, P<0.001; **, P<0.01; NS, P>0.05.

We sought to determine if loss of *hexSDF* affected resistance to other antimicrobials. We screened several other cell wall targeting antimicrobials including vancomycin, bacitracin, nisin, polymyxin, ampicillin and lysozyme. We found that loss of *hexSDF* decreased bacitracin resistance by ∼4-fold (Figure 3D), but did not result in altered resistance to other antimicrobials (Supplemental Figure 3). Expression of *hexSDF* in Δ*hexSDF* resulted in bacitracin MIC comparable to WT, however, overproduction of HexSDF in WT did not affect bacitracin MIC (Figure 3D).

We wanted to better understand the contribution of each gene in the *hexSDF* operon to daptomycin and bacitracin resistance. Thus, we constructed deletions of each gene individually. We found that loss of *hexS* had the greatest effect on daptomycin resistance, decreasing the daptomycin MIC by ∼4-fold (Figure 3E). The *hexD* and *hexF* mutations lead to a ∼2-fold decrease in the daptomycin resistance (Figure 3E). We also investigated how the individual deletions affected bacitracin resistance and found trends similar to what was found with daptomycin (Supplemental Figure 4A). We next overproduced pairs of genes from the *hex* operon and found that P_xyl_-*hexSF*, P_xyl_-*hexSD*, and P_xyl_-*hexSDF* all led to increases in daptomycin resistance (Figure 3F), while P_xyl_-*hexDF* did not lead to an increase in resistance (Figure 3F). We found that the bacitracin MICs showed a remarkably similarly trend where HexS expression is essential for increased bacitracin resistance (Supplemental Figure 4B). Combined, these data indicate that HexS is essential for increased daptomycin and bacitracin resistance.

Finally, we wanted to determine if HexSDF is important for daptomycin resistance in other strains of *C. difficile. C. difficile* 630 (PCR ribotype 012), like nearly all strains of *C. difficile*, encodes the *hex* locus (Karp et al., 2019). We utilized CRISPRi to knockdown *hexSDF* in 630Δ*erm*, and found, similarly to R20291, a ∼2-4-fold decrease in daptomycin MIC compared to the Neg sgRNA (Figure 4). This suggests that HexSDF is involved in daptomycin resistance in multiple strains of *C. difficile*.

**Figure 4.**
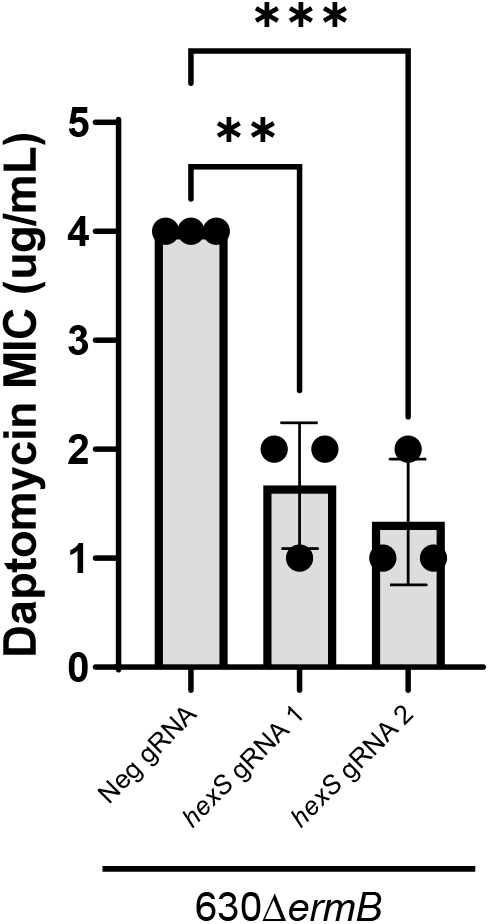
CRISPRi knockdown of *hexS* in 630Δ*erm* leads to a decrease in daptomycin MIC. Data were analyzed with one-way analysis of variance using Dunnett’s multiple comparison test. ***, P<0.001; **, P<0.01.

### HexSDF is Required for Production of HNHDRG

We were interested in the biological function of HexSDF. Based on bioinformatic predictions of the *hexSDF* operon, we hypothesized that the alterations in daptomycin and bacitracin resistance were due to changes in membrane lipid composition. Several lines of supported this: First, HexS is annotated as a monogalactosyldiacylglycerol synthase, indicating that it likely is adding sugar groups to a lipid (Altschul et al., 1990; Karp et al., 2019). Second, HexF is annotated as an MprF-like protein, MprF known in other organisms, particularly *S. aureus*, to contribute to daptomycin resistance by converting phosphatidylglycerol into lysophosphatidylglycerol (Ernst et al., 2018; Peschel et al., 2001; Pillai et al., 2007). Importantly, HexF only contains homology to the flippase domain of MprF (Supplemental Figure 2), meaning that HexF could play a role in flipping lipids across the membrane. Finally, HexD is annotated as a putative polysaccharide deacetylase, potentially removing an N-acetyl group resulting in the production of an amino-sugar. Previous work defining the polar lipids in *C. difficile* revealed the presence of a glycolipid with an amino-sugar HNHDRG (Guan et al., 2014). Based on these bioinformatic predictions, we hypothesized that HexSDF is required for production of the novel glycolipid HNHDRG.

To determine if HexSDF was required for production of glycolipids, we performed an untargeted lipidomics experiment comparing WT EV, *ΔhexSDF* EV, *ΔhexSDF* P_*xyl*_*-hexSDF, ΔhexSDF* P_*xyl*_-hexSD, *ΔhexSDF* P_*xyl*_-*hexSF*, and *ΔhexSDF* P_*xyl*_-*hexDF*. Overnight cultures were subcultured in TY supplemented with 1% xylose to an OD_600_ 0.05 and were grown to OD_600_ 0.6-0.7 at which point the cells were harvested. The membrane lipids were extracted and analyzed by HPLC-MS/MS to identify and quantify lipid species. The most striking difference we observed was the complete loss of the unique *C. difficile* glycolipid HNHDRG in the Δ*hexSDF* mutant compared to WT (Figure 5A). When HexSDF is overproduced in a Δ*hexSDF* mutant, HNHDRG is detected as similar levels to WT (Figure 5A). Interestingly, we found that the three other major glycolipids increased in Δ*hexSDF*, MHDRG (Figure 5B), DHDRG (Figure 5C), and THDRG (Figure 5D). The levels of all three glycolipids were also decreased to levels lower than WT when HexSDF was overproduced (Figure 5B, Figure 5C, Figure 5D). In contrast, phosphatidylglycerol (Figure 5E) and cardiolipin (Figure 5F) levels were not significantly different in the Δ*hexSDF* mutant or in strains overproducing HexSDF however the levels of both did trend upward in the *ΔhexSDF* mutant.

**Figure 5.**
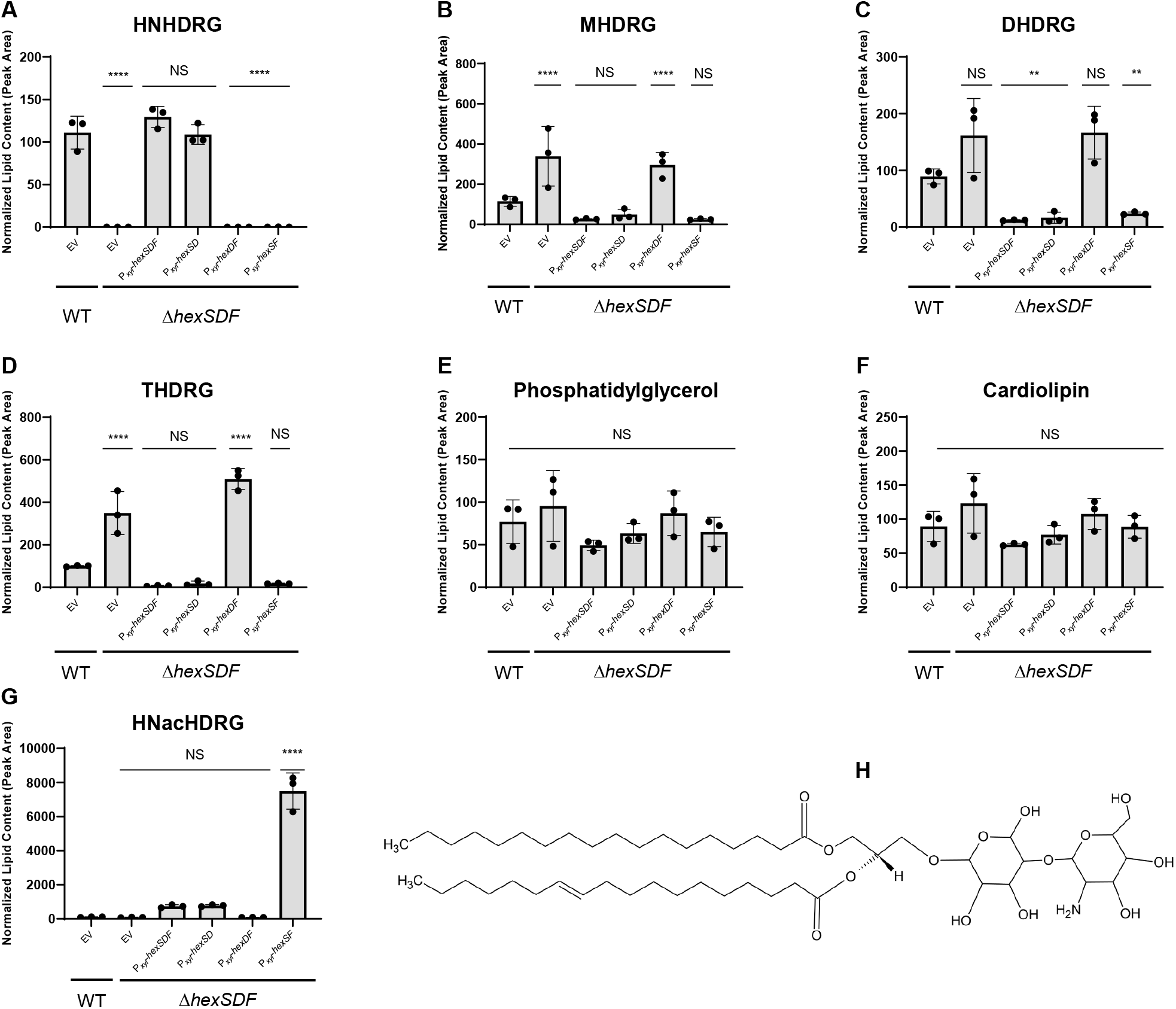
HexSDF is required for synthesis of HNHDRG. Compared to WT, Δ*hexSDF*, Δ*hexSDF* P_xyl_-*hexDF*, and Δ*hexSDF* P_xyl_-*hexSF* show significantly decreased levels of HNHDRG (A) and increased amounts of MHDRG (B), DHDRG (C), and THDRG (D). Interestingly, Δ*hexSDF* P_xyl_-*hexSF* leads to the accumulation of the HNHDRG acetylated intermediate, termed HNacHDRG (G). None of the tested strains had significant differences in phosphatidylglycerol (E) or cardiolipin (F) content. (H) The structure of HNHDRG. Data were analyzed with one-way analysis of variance using Dunnett’s multiple comparison test. ****, P<0.0001; **, P<0.01; NS, P>0.05.

We next sought to determine the contribution of individual genes in the *hex* operon. We found overproduction of HexDF did not lead to accumulation of either HNHDRG (Figure 5A). This suggests that HexS is absolutely required for HNHDRG production. Interestingly we found that overproduction of HexSF lead to accumulation of HNacHDRG, a previously undetected glycolpid which we predict is the acetylated intermediate of HNHDRG, but there was no detectable HNHDRG (Figure 5G and 5A). This suggests that the putative deacetylase HexD is required for deacetylation of HNacHDRG to produce HNHDRG. In contrast overproduction of HexSD lead to accumulation of near wild type levels of HNHDRG (Figure 5G) suggesting HexF is not essential for HNHDRG production. *in toto* we hypothesize that HexS is required for incorporation of N-acetyl-hexosyl onto the MHDRG precursor, resulting in HNacHDRG intermediate and that HexD is required for deacetylation which leads to production HNHDRG.

## Discussion

### Identification of an operon required for production of a novel glycolipid

The *C. difficile* membrane consists of ∼50% glycolipids including MHDRG, DHDRG, THDRG, and HNHDRG. We do not currently know the nature of the hexosyl groups on these lipids and in fact, we do not know if, MHDRG is a single type of glycolipid or a mixture of two or more glycolipids with different sugars. HNHDRG is a novel glycolipid that makes up ∼16% of the membrane WT *C. difficile* and to date has not been described in other bacteria (Guan et al., 2014). The principal contribution of this investigation is the identification of HexRK and HexSDF as essential for the production of the novel glycolipid HNHDRG. We identified these genes in a Tn-seq screen for genes involved in daptomycin resistance. We found that knockdowns or knockouts of either *hexRK* or *hexSDF* lead to decreases in daptomycin and bacitracin resistance. We find that in the absence of HexSDF there is almost no detectable HNHDRG. We could complement these defects with a plasmid expressing *hexSDF* from a xylose inducible promoter. This argues that HexSDF is required for resistance to daptomycin and bacitracin as well as production of the novel glycolipid HNHDRG.

### Model for HNHDRG synthesis

Based on the lipidomics data we hypothesize HNHDRG is likely produced by HexS catalyzing the addition of an N-acetylhexose to an MHDRG (Figure 6). This lipid is then flipped to the outer leaflet of the membrane by HexF, and the N-acetyl group is deacetylated by HexD, producing HNHDRG on the outer leaflet of the membrane (Figure 6). This model suggests deacetylation is required for production of HNHDRG and is supported by lipidomics data from strains lacking HexD which show the accumulation of HNacHDRG (Figure 5G). While we have identified the genes required for synthesis of HNHDRG, it is still not entirely clear what the steps of HNHDRG synthesis are. Currently, little is known about membrane lipid synthesis in *C. difficile* and if it differs compared to better studied organisms.

**Figure 6.**
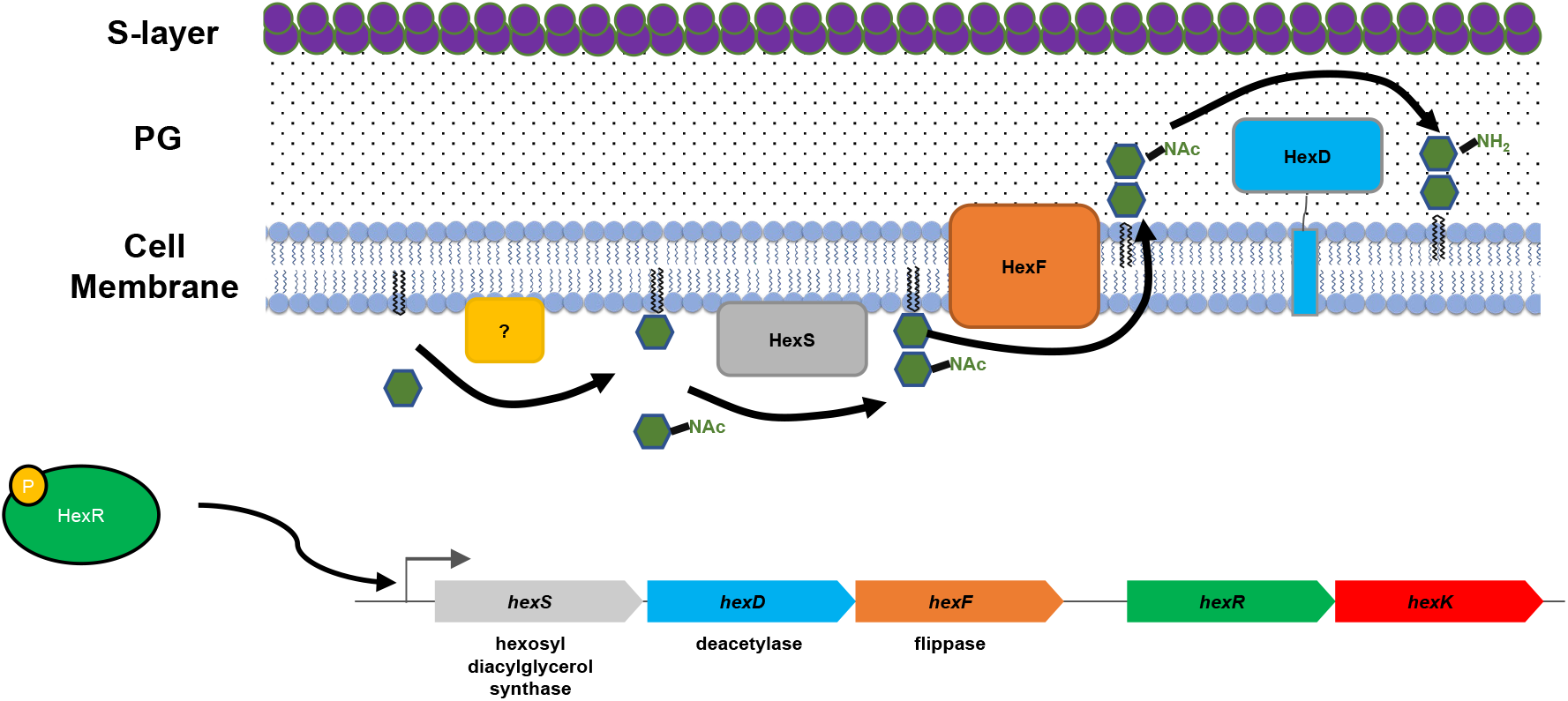
HexRK and HexSDF Model. HexR regulates the expression of *hexSDF*. We hypothesize that HexS utilizes MHDRG as its substrate and adds an N-acetyl-hexose, forming the HNacHDRG intermediate. HNacHDRG is then flipped outside the cell via HexF (and other flippases), once on the external side of the membrane NHacHDRG is deacetylated by HexD, to form HNHDRG.

In *B. subtilis* all known glycolipid synthesis is dependent upon a single enzyme UgtP which catalyzes the addition of glucose to diacylglycerol to create monoglucosyl diacylglycerol (MGDG) (Jorasch et al., 1998). In *B. subtilis* UgtP works processively to add an additional glucose group to MGDG to synthesize diglucosyl diacylglycerol (DGDG), and a third glucose to DGDG to create triglucosyl diacylglycerol TGDG (Jorasch et al., 1998). We think it is unlikely that HexS is responsible for catalyzing the addition of two different sugars, one without an N-acetyl group and one with an N-acetyl group onto diradylglycerol. Instead, we hypothesize that another enzyme, potentially one of the three *C. difficile* genes predicted to encode UgtP-like homologs (*cdr0291_0008, cdr20291_1186*, and *cdr20291_2958*) synthesizes MHDRG and HexS then adds N-acetylhexosyl to MHDRG to generate HNacHDRG. This is supported by our data showing HexS is absolutely required for HNHDRG synthesis. Consistent with this model we find increased levels of MHDRG, DHDRG and THDRG in all strains lacking HexS. We propose that in the absence of HexS MHDRG is no longer being consumed to generate HNHDRG thus more MHDRG is available to be used to synthesize DHDRG and THDRG. We also find that overproduction of HexS leads to significant decreases of MHDRG, DHDRG, and THDRG which is consistent with this model. However, at this time we cannot rule out the possibility that HexS adds Hexosyl-N-Acetylhexosyl directly to DRG (diradylglycerol).

As noted earlier we find that overproduction of HexSF without HexD results in the production of HNacHDRG but not HNHDRG. This is consistent with the role for HexD in deacetylation of HNacHDRG to produce HNHDRG. However, we were surprised that expression of HexSD without HexF produced HNHDRG at levels equivalent to all three genes alone. We hypothesize three possible explanations for this phenomenon. One possibility is that HexD functions intracellularly prior to flipping of HNHDRG. A second possibility is that HexF is not required for flipping. The third possibility is that in the absence of HexF other flippases are capable of mediating transport/flipping of HNacHDRG where it is acted on extracellularly by HexD. It is interesting to note that *C. difficile* encodes two additional genes (*cdr20291_2010* and *cdr20291_3180*) that are predicted to encode proteins with homology to HexF/MprF and may also function as flippases (Supplemental Figure 5). Taken to together these data support a model in which HexS adds N-acetylhexosyl to MHDRG leading to production of HNacHDRG. HNacHDRG is then deacetylated by HexD to produce HNHDRG.

### Identification of the HexRK regulon

HexRK represent a family of TCS that have undefined functions. We found that HexK shares homology with YrkQ in *B. subtilis*. YrkQ is a relatively unstudied TCS and *B. subtilis* is not known to produce HNHDRG nor is it known what signals activate YrkPQ. To date we have been unable to identify a signal to which HexRK responds, neither daptomycin (Figure 2C) nor bacitracin (Supplemental Figure 4C) lead to activation of the P_*hexS*_-*sLuc*^*opt*^ reporter, and our RNA-seq data suggests *hexRK* has a high basal expression (Müh et al., 2022). While the signals are not known both the HexRK and YrkPQ regulons have been identified (Kobayashi et al., 2001). Similar to the *C. difficile* HexR regulon, the YrkP regulon is small ∼8 genes and consists mostly of proteins with predicted function involved in lipid or membrane biogenesis. This includes YkcB a predicted C55-P-Glc glucosyltransferase, YkcC a predicted lipoteichoic acid glycosyltransferase, YrkO a membrane protein with 10 predicted transmembrane domains and DUF418 domain which may be involved in transport, YrkN a putative GNAT N-acetyl transferase and YrkR a 4 transmembrane domain protein with homology to phosphate-starvation-inducible protein PsiE. This raises the possibility that HexRK and YrkPQ are responsible for regulating genes involved in membrane or cell envelope biogenesis in multiple organisms. Consistent with this, the YrkP regulon is activated in daptomycin resistant mutants of *B. subtilis* (Hachmann et al., 2011). However, it is not clear how much YrkPQ contributes to daptomycin resistance in *B. subtilis*. It is also important to note that *B. subtilis* does not encode an operon homologous to *hexSDF*.

### Mechanism of HNHDRG daptomycin resistance

Since phosphatidylglycerol is required for daptomycin to kill target cells we initially expected loss of HexSDF may lead to an increase in phosphatidylglycerol content and thus a decrease in daptomycin resistance. However, we found there was not a significant change in phosphatidylglycerol levels in Δ*hexSDF* (Figure 5F). This suggests that the decrease in daptomycin resistance observed in Δ*hexSDF* is likely not due to an increase in phosphatidylglycerol levels. The subtle changes we observed in phosphatidylglycerol are still present in strains lacking either HexD or HexF. Consistent with this we find that loss of HexD or HexF also lead to a decrease in daptomycin resistance (Figure 3E). Taken together it suggests that HNHDRG is playing a more direct role in daptomycin resistance.

We are unsure how HNHDRG increases daptomycin resistance. However, one potential clue comes from the screening of the Δ*hexSDF* mutant for sensitivity to additional antibiotics where we find HexSDF is required for resistance to bacitracin and daptomycin but not other cell wall targeting antibiotics. It has been hypothesized that, since *C. difficile* does not produce PE or PS, HNHDRG serves to balance the charge of the membrane, as it can form a positively charged zwitterion (Guan et al., 2014). This could potentially explain the resistance to daptomycin and bacitracin. Both daptomycin and bacitracin require cationic metal ions to function, daptomycin requiring Ca^2+^ while bacitracin prefers Zn^2+^ though it can utilize other divalent metal ions (Adler & Snoke, 1962; Grein et al., 2020; Louie et al., 1993). We hypothesize that HNHDRG decreases the ability of positively charged metal ions to access the membrane, limiting the activity of both daptomycin and bacitracin. However, there is the caveat that loss of HNHDRG does not affect the activity of cationic antimicrobial peptides, such as lysozyme (Supplemental Figure 3), which one may expect if HNHDRG functioned by alteration of membrane charge. While the exact mechanism of resistance is unclear, we hypothesize that HNHDRG is affecting resistance directly, opposed to alterations in other membrane lipid species being responsible for resistance.

## Methods and Materials

### Bacterial strains, media, and growth conditions

Bacterial strains are listed in Table 2 The *C. difficile* strains used in this study are derivatives of R20291 (Stabler et al., 2009) and 630Δ*ermB* (Hussain et al., 2005). *C. difficile* strains were grown on tryptone-yeast (TY) medium consisted of 3% tryptone, 2% yeast extract and 2% agar (for solid medium). TY was supplemented as needed with thiamphenicol at 10 μg/mL (Thi_10_). Conjugations were performed on solid brain-heart-infusion media supplemented with yeast extract (3.65% BHI, 0.5% yeast extract, 2% agar) and plated on TY with thiamphenicol at 10 μg/mL, kanamycin at 50 μg/mL and cefoxitin at 50 μg/mL. *C. difficile* strains were maintained at 37°C in an anaerobic chamber (Coy Laboratory Products) in an atmosphere of ∼2-2.5% H_2_, 5% CO_2_, and 85% N_2_.

**Table 2.**
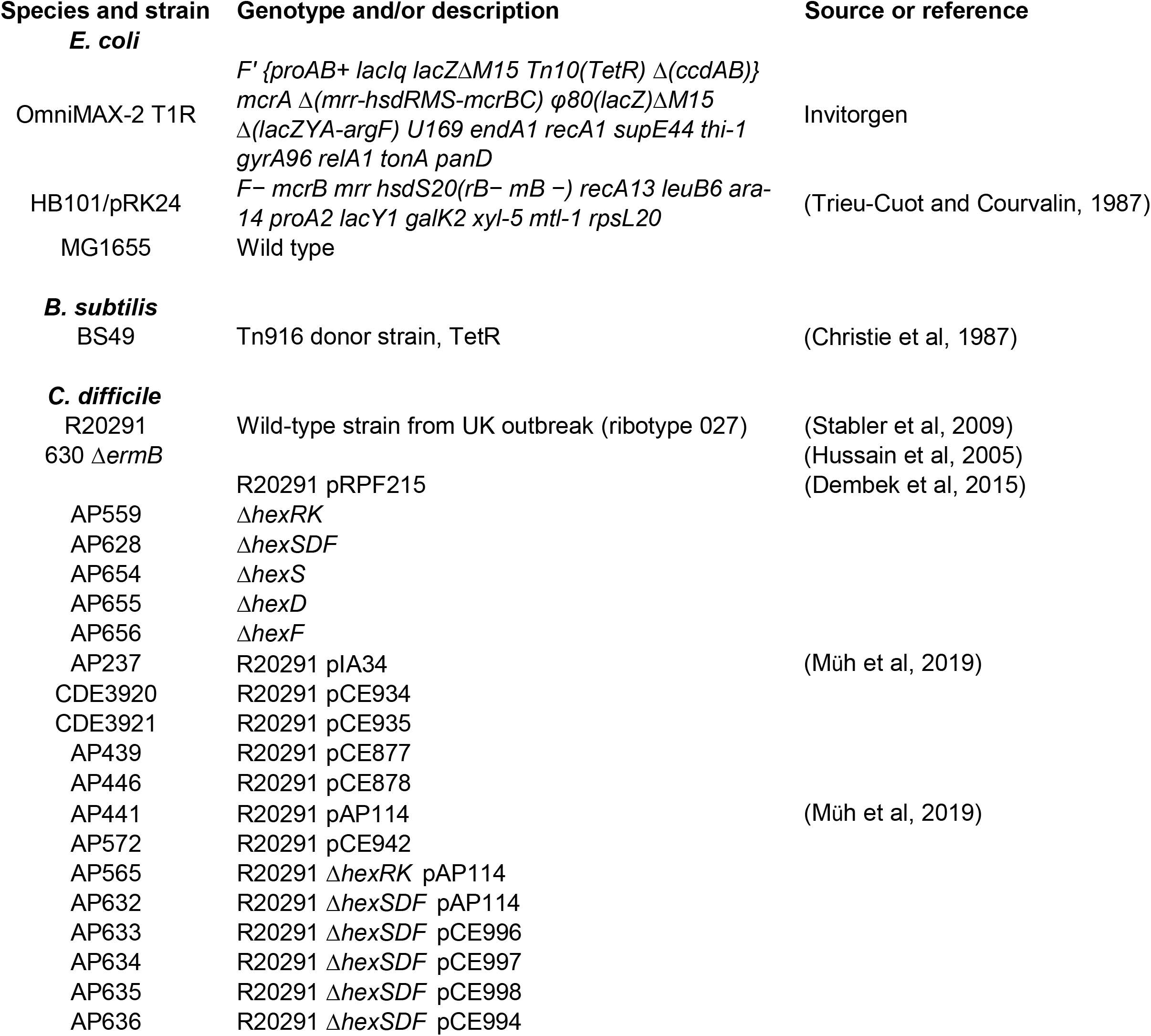
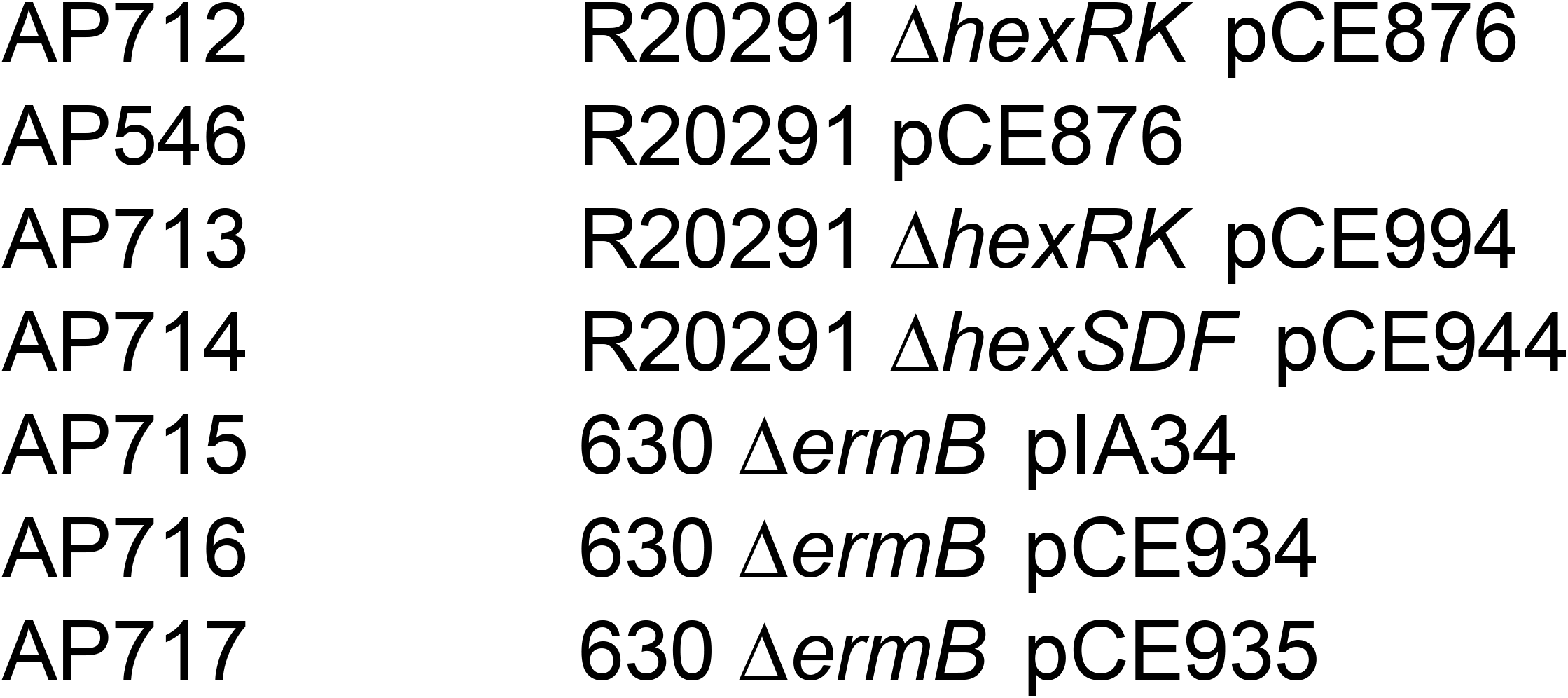
Strains.

| **Table 2 Strains** |  |  |
| --- | --- | --- |
| **Species and strain** | **Genotype and/or description** | **Source or reference** |
| ***E. coli*** |  |  |
| OmniMAX-2 T1R | *F'* {*proAB*+ *lacIq* *lacZ*Δ*M15* *Tn10*(*TetR*) Δ(*ccdAB*)}<br>*mcrA* Δ(*mrr*-*hsdRMS*-*mcrBC*) *φ80*(*lacZ*)Δ*M15*<br>Δ(*lacZYA*-*argF*) *U169* *endA1* *recA1* *supE44* *thi-1*<br>*gyrA96* *relA1* *tonA* *panD* | Invitorgen |
| HB101/pRK24 | *F*- *mcrB* *mrr* *hsdS20*(*rB*- *mB* -) *recA13* *leuB6* *ara*-<br>14 *proA2* *lacY1* *galK2* *xyl-5* *mtl-1* *rpsL20* | (Trieu-Cuot and Courvalin, 1987) |
| MG1655 | Wild type |  |
| ***B. subtilis*** |  |  |
| BS49 | Tn916 donor strain, TetR | (Christie et al, 1987) |
| ***C. difficile*** |  |  |
| R20291 | Wild-type strain from UK outbreak (ribotype 027) | (Stabler et al, 2009) |
| 630 Δ*ermB* |  | (Hussain et al, 2005) |
|  | R20291 pRPF215 | (Dembek et al, 2015) |
| AP559 | Δ*hexRK* |  |
| AP628 | Δ*hexSDF* |  |
| AP654 | Δ*hexS* |  |
| AP655 | Δ*hexD* |  |
| AP656 | Δ*hexF* |  |
| AP237 | R20291 pIA34 | (Müh et al, 2019) |
| CDE3920 | R20291 pCE934 |  |
| CDE3921 | R20291 pCE935 |  |
| AP439 | R20291 pCE877 |  |
| AP446 | R20291 pCE878 |  |
| AP441 | R20291 pAP114 | (Müh et al, 2019) |
| AP572 | R20291 pCE942 |  |
| AP565 | R20291 Δ*hexRK* pAP114 |  |
| AP632 | R20291 Δ*hexSDF* pAP114 |  |
| AP633 | R20291 Δ*hexSDF* pCE996 |  |
| AP634 | R20291 Δ*hexSDF* pCE997 |  |
| AP635 | R20291 Δ*hexSDF* pCE998 |  |
| AP636 | R20291 Δ*hexSDF* pCE994 |  |

|  |  |
| --- | --- |
| AP712 | R20291 $\Delta hexRK$ pCE876 |
| AP546 | R20291 pCE876 |
| AP713 | R20291 $\Delta hexRK$ pCE994 |
| AP714 | R20291 $\Delta hexSDF$ pCE944 |
| AP715 | 630 $\Delta ermB$ pIA34 |
| AP716 | 630 $\Delta ermB$ pCE934 |
| AP717 | 630 $\Delta ermB$ pCE935 |

*E. coli* strains were grown in LB medium (1% tryptone, 0.5% yeast extract, 0.5% NaCl, and 1.5% agar for solid medium) at 37°C with chloramphenicol at 10 μg/mL and ampicillin at 100 μg/mL as needed.

### Plasmid and bacterial strain construction

All plasmids are listed in Table 3. Plasmids were constructed using Gibson Assembly (New England Biolabs, Ipswich, MA). Regions of the plasmids constructed using PCR were verified by DNA sequencing. Oligonucleotide primers used in this work were synthesized by Integrated DNA Technologies (Coralville, IA) and are listed in Table S3. All plasmids were propagated using OmniMax-2 T1R as a cloning host. CRIPSR-Cas9 deletion plasmids were passaged through *E. coli* strain MG1655 before transformation into *B. subtilis* strain BS49 (Christie et al., 1987). CRISPR-Cas9 plasmids were built on the backbone of pJK02 (McAllister et al., 2017) with some modification (Kaus et al., 2020).

**Table 3.**
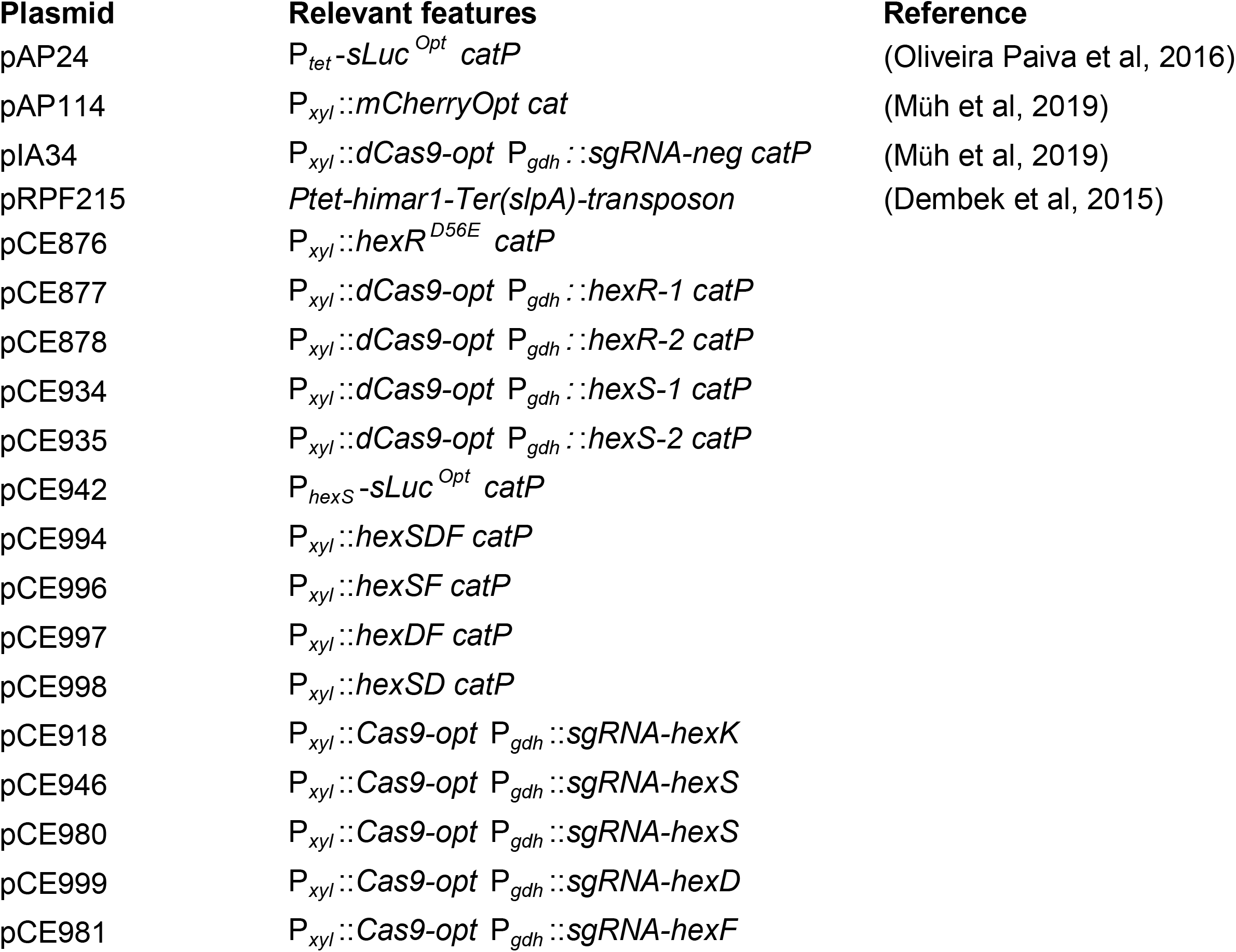
Plasmids.

For xylose-inducible overexpression constructs genes of interest were amplified using PCR, the oligonucleotides are listed in Table S3 in the supplemental material. PCR amplicons were then inserted into the plasmid pAP114 at the SacI and BamHI sites, as described previously (Müh et al., 2019). For CRISPRi constructs, two guides were created for each targeted gene of interest, guides were amplified using PCR, the oligonucleotide sequence for the guides are listed in Table S3 in the supplemental material. PCR amplicons were then inserted into the pIA33 backbone at MscI and NotI sites as previously described (Müh et al., 2019). For Nano Luciferase reporter constructs, promoters of interest were amplified using PCR, the oligonucleotides are listed in Table S3 in the supplemental material. PCR amplicons were then inserted into pAP24 (Oliveira Paiva et al., 2016) at KpnI and SacI sites using Gibson assembly. These plasmids were then passaged through *E. coli* HB101/pRK24 for conjugation into *C. difficile* (Trieu-Cuot et al., 1987)

### Antibiotic MIC determination

Overnight cultures of *C. difficile* were subcultured, grown to late log phase (OD_600_ of 1.0), and then diluted into TY to 10^6^ CFU/mL. For strains containing vectors with xylose-inducible elements, 1% xylose was added to the overnight cultures and the subcultures. A series of antibiotic concentrations was prepared in a 96-well plate in 50 μl TY broth. Wells were inoculated with 50 μL of the dilute late-log-phase culture (0.5 × 10^5^ CFU/well) and grown at 37°C for 16 hours. For strains containing vectors with xylose-inducible elements, 1% xylose was included in the media. After the incubation period the 96 well plates were spun down for 5 minutes at 5000 RPM and the MIC was determined based on the presence of cell pellets. For lysozyme MICs the steps remain the same, with the exception that after the 16 hour incubation 5 μL from each well is plated on TY for viability. The MIC from this viability plate is read the following day, with the MIC being defined as the lowest concentration of lysozyme at which 5 or fewer colonies were found per spot.

### Nano Luciferase Assays

Overnight cultures of *C. difficile* were subcultured 1:100 and grown until mid-log phase (OD_600_ ∼0.5). For antibiotic induction experiments, a series of antibiotic concentrations was prepared in a 96-well plate in 50 μL TY broth. Wells were inoculated with 50 μL of mid-log phase culture. Inoculated plates were allowed to incubate for 2 hours at 37°C in the anaerobic chamber. 50 μL from each well was then transferred to solid white 96-well plates. A 1:100 dilution was made of the Nano-Glo Luciferase Assay (Promega) working solution. 50 μL of the diluted Nano-Glo solution was added to each well in the white plate. Plates were allowed to rest at room temperature for 5 minutes prior to luminescence measurement. Luminescence was measured using an Infinite M200 Pro plate reader (Tecan). Luminescence values were normalized to OD_600_.

### RNA Extraction

Overnight cultures were grown in TY then subcultured 1:100 in 100 mL of TY. Subcultures were grown to OD_600_ 0.7-0.8, mixed 1:1 with a solution of 50% ethanol 50% acetone and immediately frozen at -80 °C. Samples were stored at -80 °C until ready for harvest. Extraction of RNA was performed using the RNeasy extraction kit (Qiagen) as previously described (Müh et al., 2022). RIN was determined using the Agilent BioAnalyzer 2000 at the University of Iowa.

### RNA-seq and Analysis

Sequencing was performed by (SeqCenter, Pittsburgh, PA) using 51 bp paired end reads on. We achieved a read depth of ∼80-120X per sample. Data analysis was done using the ProkSeq (Mahmud et al., 2021) pipeline, which performs trimming and QC using, genome alignment using Bowtie2, quantification of RNA expression levels using FeatureCounts, and differential expression analysis using DEseq2 (Love et al., 2014). The full RNA-seq dataset can be found in Table S2.

### Lipidomics

Overnight cultures were grown in TY supplemented with 1% xylose and 10 μg/mL thiamphenicol in biological triplicate. Overnight cultures were then subcultured to an OD_600_ 0.05 in TY supplemented with 1% xylose and 10 μg/mL thiamphenicol. Subcultures were allowed to grow until OD_600_ 0.6-0.7 at which point the cells were harvested and pelleted at 8000 rpm for 20 minutes. Biological replicates (3) were grown on different days.

Lipid extraction and LC-MS/MS was performed by Cayman Chemical Company. After thawing, cells were mixed with 5 mL methanol, transferred to 7 mL Precellys tubes containing 0.1mm ceramic beads (Bertin Technologies, CK01 lysing Kit) and homogenized with 3 cycles at 8800 rpm, for 30s, with 60 s pauses between cycles. Then, 800 μL of the homogenized mixtures were transferred to 8 mL screw-cap glass tubes. A methyl-tert-butyl ether (MTBE)-based liquid-liquid extraction protocol was used, by first adding 1.2 mL methanol containing a mixture of deuterated internal standards covering several major lipid categories (fatty acids, glycerolipids, glycerophospholipids, sphingolipids, and sterols), and then 4 mL MTBE. The mixture was incubated on a tabletop shaker at 500 rpm at room temperature for 1 h, and then stored at 4 °C for 60 h to maximize lipid extraction. After bringing the samples to room temperature, phase separation was induced by adding 1 mL water to each sample. The samples were vortexed and then centrifuged at 2000 x *g* for 15 min. The upper organic phase of each sample was carefully removed using a Pasteur pipette and transferred into a clean glass tube. The remaining aqueous phase was reextracted with 2 mL of the upper phase of MTBE/methanol/water 10:3:2.5 (v/v/v). After vortexing and centrifuging, the organic phase was collected and combined with the initial organic phase. The extracted lipids were dried overnight in a SpeedVac vacuum concentrator.

The dried lipid extracts were reconstituted in 200 μL n-butanol/methanol 1:1 (v/v)and transferred into autosampler vials for analysis by LC-MS/MS. Aliquots of 5 μL were injected into an Ultimate 3000 UPLC system connected to a Q Exactive Plus Orbitrap mass spectrometer (Thermo Scientific). An Accucore C30 2.6 μm, 150 × 2.1 mm HPLC column (Thermo Scientific) was used, using mobile phases A, (acetonitrile/water/formic acid 60:40:0.1 (v/v/v), containing 10 mM ammonium formate) and B, (acetonitrile/isopropanol/formic acid 10:90:0.1 (v/v/v), containing 10 mM ammonium formate). Lipids were eluted at a constant flow rate of 300 μL/min using a gradient from 30% to 99% mobile phase B over 30 min. The column temperature was kept at a constant 40 °C. Polarity switching was used throughout the gradient to acquire high-resolution MS data (resolution 75,000) and data-dependent MS/MS data.

Data analysis was performed using Lipostar software (Version 2; Molecular Discovery) for detection of features (peaks with unique *m/z* and retention time), noise and artifact reduction, alignment, normalization, and lipid identification. Automated lipid identification was performed by querying the Lipid Maps Structural Database (LMSD), modified by Cayman to include many additional lipids not present in the LMSD.

Statistical analysis of the data presented on Figure 5 was performed on the summed peak areas of lipids with the same head groups, normalized to the WT values to allow for comparison between strains. Data were analyzed with one-way analysis of variance using Dunnett’s multiple comparison test.

### Transposon Mutagenesis of *C. difficile*

We generated a pooled transposon library in R20291 consisting of ∼80,000 colonies using pRPF215 as previously described (Dembek et al., 2015). Briefly, mid-log *C*. difficile R20291 containing pRPF215 was plated on TY supplemented with 20 μg/mL lincomycin and 50 ng/mL of anhydrotetracycline. After overnight growth we pooled the library. To identify genes required for daptomycin resistance we grew the pooled transposon library to with or without 2 μg/mL daptomycin. Briefly, 100 μL of the prepared transposon library was subcultured into 5 mL of TY supplemented with 20 μg/mL lincomycin and 100 ng/mL anhydrotetracycline. Subcultures were grown until OD_600_ 0.2-0.3 at which point 0 or 2 μg/mL daptomycin was added. Cultures were incubated with daptomycin until OD_600_ 1.0, cells were then pelleted. gDNA was extracted by washing cell pellets 3 times in 1X PBS. Pellets were suspended in 300 μL of 1X thermopol buffer (New England Biolabs) and 5 μL of 800 U/mL proteinase K (New England Biolabs) was added. Cells were incubated in solution at 37 °C for 1 hour until they were clearly lysed. The DNA was extracted using Phenol:Chloroform:Isoamyl Alcohol (25:24:1, Fischer Scientific Company) followed by Ethanol precipitation of the DNA. The purified gDNA was resuspended in 100 μL of ddH_2_0.

### Sequencing Analysis of Transposon Insertions

Preparation of the transposon amplicon library was performed as previously described (Karash et al., 2019). Briefly the primer CDEP4815 was used to perform multiple rounds of single primer extension. The extension products were then C-tailed using terminal transferase (New England Biolabs). The resulting C-tailed extension products were then amplified using primers CDEP4816 and P1 or CDEP4816 and P3 to generate barcoded amplicon libraries for Illumina sequencing. Fragments of 300-500 bp were extracted from a gel and purified. Sequencing was performed on Illumina HiSeq X with 150bp PE reads. Sequencing was performed by Admera Health (South Plainfield, NJ).

The resulting read data were analyzed using the Transit package for TnSeq analysis (DeJesus et al., 2015). Briefly the reads were demultiplexed using Sabre (Najoshi, 2011). The data was then trimmed and aligned to eliminate poor quality reads using the Transit Pre-processor (DeJesus et al., 2015). Transit aligned the trimmed reads to the *C. diffiicle* R20291 genome using BWA, tabulated Tn insertion counts, and calculated conditional essentially utilizing the resampling comparison method (DeJesus et al., 2015)

## Data availability

RNA-seq data were submitted to the NCBI GEO repository and assigned accession number GSE220119.

## Acknowledgements

This work was supported by Public Health Service grant R01AI087834 to CDE from the National Institutes for Allergy and Infectious Disease. AP was supported by T32AI007511. We thank David Weiss and members of the Ellermeier lab for helpful discussions.

## Figure Legends

**Supplemental Figure 1**. Protein alignment of HexK and YrkQ made using ClustalW.

**Supplemental Figure 2**. Protein alignment of HexF and *B. subtilis* MprF made using ClustalW. The flippase domain of *B. subtilis* MprF has been highlighted in red.

**Supplemental Figure 3**. Δ*hexSDF* MICs of other antimicrobials including: (A). vancomycin (B). Nisin (C). Polymyxin B (D). Ampicillin (E). Lysozyme.

**Supplemental Figure 4**. The *hex* locus mediates resistance to bacitracin. (A) Δ*hexS* and Δ*hexD* led to significant decreases in bacitracin MIC compared to WT, however, Δ*hexF* did not have a significant decrease in bacitracin MIC. (B) Bacitracin MIC comparing overproduction of HexSF, HexSD, and HexDF in Δ*hexSDF*. Overproduction of HexSF and HexSD led to mild increases in bacitracin resistance, while overproduction of HexDF did not lead to notable increases in bacitracin resistance. (C) P_*hexS*_-*sLuc*^*opt*^ did not show altered luminescence when exposed to varying concentrations of bacitracin for 2 hours.

**Supplemental Figure 5**. Protein multiple sequence alignment comparing HexF, CDR2010, and CDR3180 made using ClustalW.

